# Testing the effectiveness of principal components in adjusting for relatedness in genetic association studies

**DOI:** 10.1101/858399

**Authors:** Yiqi Yao, Alejandro Ochoa

## Abstract

Modern genetic association studies require modeling population structure and family relatedness in order to calculate correct statistics. Principal Components Analysis (PCA) is one of the most common approaches for modeling this population structure, but nowadays the Linear Mixed-Effects Model (LMM) is believed by many to be a superior model. Remarkably, previous comparisons have been limited by testing PCA without varying the number of principal components (PCs), by simulating unrealistically simple population structures, and by not always measuring both type-I error control and predictive power. In this work, we thoroughly evaluate PCA with varying number of PCs alongside LMM in various realistic scenarios, including admixture together with family structure, measuring both null p-value uniformity and the area under the precision-recall curves. We find that PCA performs as well as LMM when enough PCs are used and the sample size is large, and find a remarkable robustness to extreme number of PCs. However, we notice decreased performance for PCA relative to LMM when sample sizes are small and when there is family structure, although LMM performance is highly variable. Altogether, our work suggests that PCA is a favorable approach for association studies when sample sizes are large and no close relatives exist in the data, and a hybrid approach of LMM with PCs may be the best of both worlds.

## 1 Introduction

The goal of a genetic association study is to identify loci whose genotypes are correlated significantly with a certain trait. An important assumption made by classical association tests is that genotypes are unstructured: drawn independently from a common allele frequency. However, this assumption does not hold for structured populations, which includes multiethnic cohorts and admixed individuals, and for family data. When naive approaches are incorrectly applied to structured populations and/or family data, association statistics (such as *χ*^2^) become inflated relative to the null expectation, resulting in greater numbers of false positives than expected and loss of power (Devlin and Roeder, 1999; Voight and Pritchard, 2005; Astle and Balding, 2009).

The most popular approaches for conducting genetic association studies with structured populations involve modeling the population structure via covariates. Such covariates may be inferred ancestry proportions (Pritchard et al., 2000) or transformations of these. Principal components analysis (PCA) represents the most common of these variants, in which the top eigenvectors of the kinship matrix are used to model the population structure (Zhang et al., 2003; Price et al., 2006; Bouaziz et al., 2011). These top eigenvectors are commonly referred to as Principal Components (PCs) in the genetics literature (the convention we adopt here; Patterson et al., 2006), but it is worth noting that in other fields the PCs would instead denote the projections of the data onto the eigenvectors (Jolliffe, 2002). Various works have found that PCs map to ancestry, and PCs work as well as ancestry in association studies and can be inferred more quickly (Patterson et al., 2006; Zhao et al., 2007; Bouaziz et al., 2011). More recent work has focused on speeding up the calculation of PCs rather than on evaluating its performance in association studies (Lee et al., 2012; Abraham and Inouye, 2014; Galinsky et al., 2016; Abraham et al., 2017). PCA remains a popular and powerful approach for association studies (Wojcik et al., 2019).

The other dominant approach for genetic association studies under population structure is the Linear Mixed-effect Model (LMM), in which population structure is a random effect drawn from a covariance model parametrized by the kinship matrix. LMM and PCA share deep connections that suggest that both models ought to perform similarly (Astle and Balding, 2009; Janss et al., 2012; Hoffman, 2013). However, many previous studies have found that LMM outperforms the PCA approach, although many evaluations have been limited (Zhao et al., 2007; Astle and Balding, 2009; Kang et al., 2010). Other studies find that PCA can outperform LMM in certain settings (Price et al., 2010; Wu et al., 2011; Wang et al., 2013), although these are believed to be unusual (Sul and Eskin, 2013). Moreover, various explanations for if and why LMM outperforms PCA are vague and have not been tested directly (Price et al., 2010; Sul and Eskin, 2013; Price et al., 2013; Hoffman, 2013). Since LMMs tend to be considerably slower than the PCA approach, it is important to understand when the difference in performance between these two approaches is outweighed by their difference in runtime.

PCA has been evaluated in numerous previous works in the context of association studies. However, all of these studies have important limitations, for the most part due to PCA being treated as a competitor rather than a method worthy of exploring more fully. For example, although there are methods for selecting the numbers of PCs (Patterson et al., 2006), most evaluations either admit to selecting 10 because it has long been the default and it performs well enough, regardless of the dataset in question (Epstein et al., 2007; Li and Yu, 2008; Astle and Balding, 2009; Li et al., 2010; Wu et al., 2011), or test only one number of PCs without much justification (Zhang et al., 2003; Kimmel et al., 2007; Zhao et al., 2007; Zhang et al., 2008; Price et al., 2010; Bouaziz et al., 2011; Hoffman, 2013; Wang et al., 2013; Tucker et al., 2014; Yang et al., 2014; Sul et al., 2018). Conversely, only a few studies consider a (small) set of numbers of PCs, where they show remarkable robustness to this choice (Price et al., 2006; Kang et al., 2010; Wojcik et al., 2019). Moreover, most of these evaluations considered simulated data with only *K* = 2 independent subpopulations or admixture from only two subpopulations (exceptions are Astle and Balding (2009) with *K* = 3, and Wang et al. (2013) with *K* = 4), although worldwide human population structure is expected to have a larger dimensionality of at least *K* = 9 (Wojcik et al., 2019). Similarly, only one evaluation simulated data from a family pedigree: Price et al. (2010) included sibling pairs. Some studies did include evaluations involving real data that featured known or cryptic relatedness, but thes analyses did not measure type-I error rates or power calculations, most of which settled for measuring test statistic inflation. Lastly, many of the earlier evaluations employed case-control simulations exclusively (as opposed to quantitative traits as we do here), were based on very small real or simulated datasets relative to today’s standards, did not include any LMMs in their evaluations, and often did not measure both type-I error rates and power (or one of their proxies). Here we aim to systematically evaluate the robustness of the PCA approach to the choice of number of PCs, especially in cases where the model is grossly misspecified, and to compare to the gold standard LMM approach in more realistic simulations relevant to today’s research.

In this work, we study the performance of the PCA method in genetic association studies, comparing it to a leading LMM approach, characterizing its behavior under various numbers of PCs and varying sample sizes, under a reasonable admixture model with *K* = 10 source subpopulations and also a model with admixture and family structure. Our evaluation is more thorough than previous ones, directly measuring the uniformity of null p-values (as required for accurate type-I error control and FDR control via q-values; Storey, 2003; Storey and Tibshirani, 2003) and predictive power by calculating the area under precision-recall curves. We find that the performance of PCA is favorable when sample sizes are large (at least 1,000 individuals), matching the performance of LMMs as long as enough PCs are used. Remarkably, the approach is robust even when the number of PCs far exceeds the optimal number for reasonably large studies. However, for smaller studies (100 individuals) there is a more pronounced loss of power when the number of PCs exceeds the optimal number. Moreover, LMMs outperform PCA in our small sample size simulation and in the presence of family structure, which is a well-known case where PCA fails (Patterson et al., 2006; Price et al., 2010). All together, our simulation studies provide clear criteria under which use of PCA results in acceptable performance compared to LMMs.

## 2 Models and Methods

### 2.1 Models for genetic association studies

In this subsection we describe the complex trait model and kinship model that motivates both the PCA and LMM models for genetic association studies, followed by further details regarding the PCA and LMM approaches. The derivations of the PCA and LMM models from the general quantitative trait model are similar to previous presentations (Astle and Balding, 2009; Janss et al., 2012; Hoffman, 2013), but we emphasize the kinship model for random genotypes as being crucial for these connections, and make a clear distinction between the true kinship matrix and its most common estimator, which is biased (Ochoa and Storey, 2016b; Ochoa and Storey, 2018).

#### 2.1.1 The complex trait model and PCA approximation

Let *x*_*ij*_ ∈ {0, 1, 2} be the genotype at locus *i* for individual *j*, which counts the number of reference alleles. Suppose there are *n* individuals and *m* loci, **X** = (*x*_*ij*_) is their *m* × *n* genotype matrix, and **y** is the length-*n* (column) vector which represents trait value for each individual. The approaches we consider are based on the following additive linear model for a quantitative (continuous) trait:

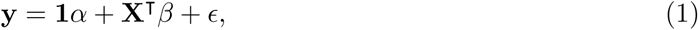

where **1** is a length-*n* vector of ones, *α* is the scalar intercept coefficient, *β* is the length-*m* vector of locus effect sizes, and *ϵ* is a length-*n* vector of residuals. The residuals are assumed to follow a normal distribution: *ϵ*_*j*_ ∼ Normal(0, *σ*^2^) independently for each individual *j*, for some residual variance parameter *σ*^2^.

Typically the number of loci *m* is in the order of millions while the number of individuals *n* is in the thousands. Hence, the full model above cannot be fit in this typical *n* ≪ *m* case, as there are only *n* datapoints to fit (the trait vector) but there are *m* + 1 parameters to fit (*α* and the *β* vector). The PCA model with *r* PCs corresponds to the following approximation to the full model, corresponding to a model fit at a single locus *i*:

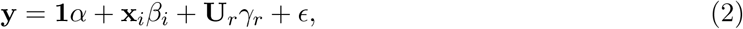

where **x**_*i*_ is the length-*n* vector of genotypes at locus *i* only, *β*_*i*_ is the effect size coefficient for that locus, **U**_*r*_ is an *n* × *r* matrix of PCs, and *γ*_*r*_ is the length-*r* vector of coefficients for the PCs. This approximation is explained by first noticing that the genotype matrix has the following singular value decomposition: **X**^**┬**^ = **UDV**^**┬**^, where assuming *n < m* we have that **U** is an *n* × *n* matrix of the left singular vectors of **X, V** is an *m* × *n* matrix of its right singular vectors, and **D** is an *n* × *n* diagonal matrix of its singular values. Thus, in the full model we have **X**^**┬**^*β* = **U***γ*, where *γ* = **DV**^**┬**^*β* is a length-*n* vector. The approximation consists solely of replacing **U***γ* (the full set of *n* left singular vectors and their coefficients) with **U**_*r*_*γ*_*r*_ (the top *r* singular vectors only, which constitutes the best approximation of rank *r*). Thus, the extra terms in the PCA approach approximate the polygenic effect of the whole genome, and assumes that the locus *i* being tested does not contribute greatly to this signal.

The statistical significance of a given association test is performed as follows. The null hypothesis is *β*_*j*_ = 0 (no association). The null and alternative models are each fit (fitting the coefficients of the multiple regression, where *β*_*j*_ is excluded under the null while it is fit under the alternative). The resulting regression residuals are compared to each other using the F-test, which results in a two-sided p-value. Note that many common PCA implementations trade the more exact F-test for a *χ*^2^ test, which is simpler to implement but only asymptotically accurate. As this is a multiple hypothesis test, there are a large number of loci (*m*) tested for association, so it is best to control the FDR rather than setting a fixed p-value threshold. We recommend estimating q-values and setting a threshold of *q <* 0.05 so that the FDR is controlled at the 5% level.

#### 2.1.2 Kinship model for genotypes

In order to better motivate the most common estimation procedure of PCs for genotype data, and to connect PCA to LMMs, we shall review the kinship model for genotypes. The model states that genotypes are random variables with a mean and covariance structure given by

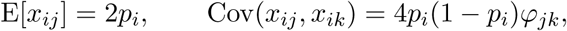

where *p*_*i*_ is the ancestral allele frequency at locus *i* and *ϕ*_*jk*_ is the kinship coefficient between individuals *j* and *k* (Malécot, 1948; Wright, 1951; Jacquard, 1970). Thus, if we standardize the genotype matrix as

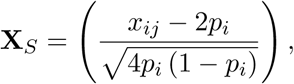

then this results in a straightforward kinship matrix estimator:

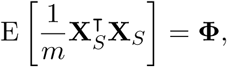

where **Φ** = (*ϕ*_*jk*_) is the *n* × *n* kinship matrix. Note that replacing the raw genotype matrix **X** with the standardized matrix **X**_*S*_ in the trait model of Eq. (1) results in an equivalent model, as this covariate differs only by a linear transformation. Thus, under the standardized genotype model, the PCs of interest are equal in expectation to the top eigenvectors of the kinship matrix.

#### 2.1.3 Estimation of principal components from genotype data

In practice, the matrix of principal components **U**_*r*_ in Eq. (2) is determined from an estimate of the earlier standardized genotype matrix **X**_*S*_, namely

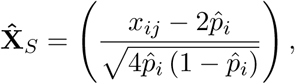

where the true ancestral allele frequency *p*_*i*_ is replaced by the estimate 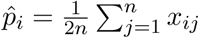, and results in the kinship estimate 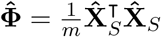 This kinship estimate and minor variants are also employed in LMMs (Yang et al., 2011). This estimator of the kinship matrix is biased, and this bias is different for every individual pair (Ochoa and Storey, 2016b; Ochoa and Storey, 2018). However, in the present context of PCA regression in genetic association studies, the existing approach performs as well as when the above estimate is replaced by the true kinship matrix (not shown). Thus, it appears that in combination with the intercept term (**1***α* in Eq. (2)), the rowspace of this kinship matrix estimate approximately equals that of the true kinship matrix.

#### 2.1.4 Linear mixed-effects model

The LMM is another approximation to the complex trait model in Eq. (1), given by

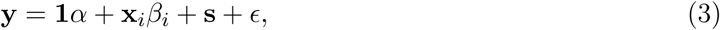

which is like the PCA model in Eq. (2) except that the PC terms **U**_*r*_*γ*_*r*_ are replaced by the random effect **s**, which is a length-*n* vector drawn from

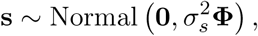

where **Φ** is the kinship matrix and 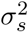 is a trait-specific variance scaling factor. This model is derived from treating the standardized genotype matrix **X**_*S*_ as random rather than fixed, so that the standardized genetic effect 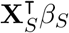 in Eq. (1) has mean zero and a covariance matrix of

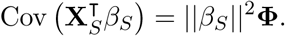

The above random effect **s** satisfies those equations, where the variance scale equals 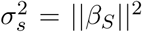. Thus, the PCA approach is the fixed model equivalent of the LMM under the additional approximation that only the top *r* eigenvectors are used in PCA whereas the LMM uses all eigenvectors.

A key advantage of LMM over PCA is that it has fewer parameters to fit: ignoring the shared terms in Eq. (2) and Eq. (3), PCA has *r* parameters to fit (each PC coefficient in the *γ* vector), whereas LMMs only fit one additional parameter, namely 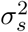. Therefore, PCA is expected to overfit more substantially than LMM—and thus lose power—when *r* is very large, and especially when the sample size (the number of individuals *n*) is very small.

Due to its accuracy and speed, the LMM implementation that we chose for our evaluations is GCTA (Yang et al., 2011). GCTA uses the same biased kinship matrix estimator 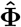 as standard PCA approaches, and the version that incorporates PCs also derives the PCs from the same kinship matrix estimate; thus, when both kinship and PCs are used, the population structure is essentially modeled twice, although previous work has found this apparent redundancy beneficial (Zhao et al., 2007; Price et al., 2010). It is worth noting that earlier LMM approaches estimated kinship matrices using maximum likelihood approaches that excluded population structure from their estimates, and population structure was modeled via admixture proportions rather than PCA (Yu et al., 2006; Zhao et al., 2007).

### 2.2 Simulations

#### 2.2.1 Genotype simulation from the admixture model

We consider three simulation scenarios, refered to as (1) large sample size, (2) small sample size, and (3) family structure. All cases are based on the admixture model described previously (Ochoa and Storey, 2016a; Ochoa and Storey, 2016b), and which is implemented in the R package bnpsd available on GitHub and the Comprehensive R Archive Network (CRAN).

Here we consider scenarios where the number of individuals *n* varies: the large sample size and family structure scenarios have *n* = 1, 000 whereas small sample size has *n* = 100. The number of loci in all cases is *m* = 100, 000. Individuals are admixed from *K* = 10 intermediate subpopulations, where *K* is also the rank of the population structure; thus, after taking into account the intercept’s rank-1 contribution, the population structure can be fit with *r* = *K* − 1 PCs. Each subpopulation *S*_*u*_(*u* ∈ {1, …, *K*}) has an inbreeding coefficient *f*_*S*_*u* = *uτ*, individual-specific admixture proportions *q*_*ju*_ for individual *j* and intermediate subpopulation *S*_*u*_ arise from a random walk model for the intermediate subpopulations on a 1-dimensional geography with spread *σ*, where the free parameters *τ* and *σ* are fit to result in *F*_ST_ = 0.1 for the admixed individuals and a bias coefficient of *s* = 0.5, exactly as before (Ochoa and Storey, 2016b).

Random genotypes are drawn from this model, as follows. First, uniform ancestral allele frequencies *p*_*i*_ are drawn. The allele frequency 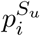 at locus *i* of each intermediate subpopulation *S*_*u*_ is drawn from the Beta distribution with mean *p*_*i*_ and variance *p*_*i*_(1 − *p*_*i*_)*f*_*S*_*u* (Balding and Nichols, 1995). The individual-specific allele frequency of individual *j* and locus *i* is given by 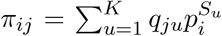.Lastly, genotypes are drawn from *x*_*ij*_ ∼ Binomial(2, *π*_*ij*_). Loci that are fixed (where for some *i* we had *x*_*ij*_ = 0 for all *j*, or *x*_*ij*_ = 2 for all *j*) are drawn again from the model, starting from *p*_*i*_, iterating until no loci are fixed.

#### 2.2.2 Genotype simulation from the family model

Here we describe a simulation of a family structure with admixture that aims to be realistic by: (1) pairing all individuals in every generation, resulting in two children per couple; (2) strictly avoiding close relatives when pairing individuals; (3) strongly favoring pairs that are nearby in their 1-dimensional geography, which helps preserve the population structure across the generations by preferentially pairing individuals with more similar admixture proportions (a form of assortative mating); and (4) iterating for many generations so that a broad distribution of close and distant relatives is present in the data.

Generation 1 has individuals with genotypes drawn from the large sample size scenario described earlier, which features admixture. In subsequent generations, every individual is paired as follows. The local kinship matrix of individuals is stored and updated after every generation, which records the pedigree relatedness; in the first generation, everybody is locally unrelated. Also, individuals are ordered, initially by the 1-dimensional geography, and in subsequent generations paired individuals are grouped and reordered by their average coordinate, preserving the original order when there are ties. For every remaining unpaired individual, one is drawn randomly from the population, and it is paired with the nearest individual that is not a second cousin or closer relative (local kinship must be *<* 1*/*4^3^). Note that every individual is initially genderless, and after pairing one individual in the pair may be set to male and the other to female without giving rise to contradictions. If there are individuals that could not be paired (occurs if unpaired individuals are all close relatives), then the process of pairing individuals randomly is repeated entirely for this generation. If after 100 iterations no solution could be found randomly (there were always unpaired individuals), then the simulation restarts from the very first generation; this may occur for very small populations, but was not observed when *n* = 1000. Once individuals are paired, two children per pair have their genotypes drawn independently of each other. In particular, at every locus, one allele is drawn randomly from one of the parents and the other allele from the other parent. Loci are constructed independently of the rest (no linkage disequilibrium). The simulation continues for 20 generations. As this simulation is very computationally expensive, it was run only once (genotypes did not change as new random traits were constructed as described next).

#### 2.2.3 Trait Simulation

For a given genotype matrix (simulated or real), a simulated complex trait that follows the additive quantitative trait model in Eq. (1) is constructed as follows. In all cases we set the heritability of the trait to be *h*^2^ = 0.8. We varied the number of causal loci (*m*_1_) together with the number of individuals (*n*) so power would remain balanced: for the *n* = 1, 000 cases we set *m*_1_ = 100, whereas the *n* = 100 simulation had *m*_1_ = 10.

Each simulation replicate consists of different causal loci with different effect sizes, as follows. The non-genetic effects are drawn from *ϵ*_*j*_ ∼ Normal(0, 1 − *h*^2^) independently for each individual *j*. A subset of size *m*_1_ of loci was selected at random from the genotype matrix to be causal loci. The effect size *β*_*i*_ at each causal locus *i* is drawn initially from a Standard Normal distribution. At non-causal loci *i* we have *β*_*i*_ = 0. Under the kinship model, the resulting genetic variance component is given by

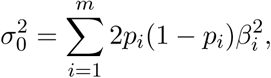

where *p*_*i*_ is the true ancestral allele frequency at locus *i*, which is known in our simulations. The desired genetic variance of *h*^2^ is therefore obtained by multiplying every *β*_*i*_ by 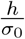. Lastly, the intercept coefficient in Eq. (1) is set to 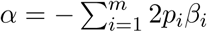, so the trait expectation is zero. This trait simulation procedure is implemented in the simtrait R package, available at https://github.com/OchoaLab/simtrait.

### 2.3 Evaluation of performance

All of the approaches considered here are evaluated in two orthogonal dimensions. The first one— the RMSD_*p*_ statistic below—quantifies the extent to which null p-values are uniform, which is a prerequisite for accurate control of the type-I error and successful FDR control via q-values. The second measure—the area under the precision-recall curve—quantifies the predictive power of each method, which makes it possible to qualitatively compare the statistical power of each method without having to select a single threshold, and most importantly, overcoming the problem of comparing methods that may not have accurate p-values (Bouaziz et al., 2011).

#### 2.3.1 RMSD_*p*_: a measure of p-value uniformity

From their definition, correct p-values (for continuous test statistics) have a uniform distribution when the null hypothesis holds. This fact is crucial for accurate control of the type-I error, and is a prerequisite for the most common approaches that control the FDR, such as q-values (Storey, 2003; Storey and Tibshirani, 2003). We use the Root Mean Square Deviation (RMSD) to measure the disagreement between the observed p-value quantiles and the expected uniform quantiles:

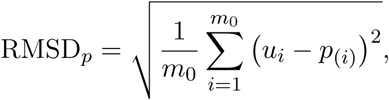

where *m*_0_ = *m* − *m*_1_ is the number of null loci (*β*_*i*_ = 0 cases only), here *i* indexes null loci only, *p*(*i*) is the *i*th ordered null p-value, and *u*_*i*_ = (*i* − 0.5)*/m*_0_ is its expectation. Thus, RMSD_*p*_ = 0 corresponds to the best performance in this test, and larger RMSD_*p*_ values correspond to worse performance.

In previous evaluations, test statistic inflation has been used to measure the success of corrections for population structure (Astle and Balding, 2009; Price et al., 2010). The inflation factor *λ* is defined as the median *χ*^2^ association statistic divided by theoretical median under the null hypothesis (Devlin and Roeder, 1999). Hence, when null test statistics have their expected distribution, we get *λ* = 1 (same as RMSD_*p*_ = 0 above). However, any other null test statistic distribution with the same median results in *λ* = 1 as well, which is a flaw of this test that RMSD_*p*_ overcomes (RMSD_*p*_ = 0 if and only if null test statistics have their expected distribution). The *λ >* 1 case (gives RMSD_*p*_ *>* 0) corresponds to inflated statistics, which occurs when residual population structure is present. *λ <* 1 is not expected for genetic association studies (also gives RMSD_*p*_ *>* 0). Note that *λ* only use the median of the null distribution, whereas the RMSD_*p*_ makes use of the complete p-value distribution to evaluate its uniformity, which is more accurate. The drawback is that RMSD_*p*_ requires knowing which loci are null, so unlike *λ*, it is not applicable to the p-values of real association studies.

#### 2.3.2 The area under the precision-recall curve

Precision and recall are two common measures for evaluating binary classifiers. Let *c*_*i*_ be the the true classification of locus *i*, where *c*_*i*_ = 1 for truly causal loci (if the true *β*_*i*_ ≠ 0, where the alternative hypothesis holds), and *c*_*i*_ = 0 otherwise (null cases). For a given method and some threshold *t* on its per-locus test statistics, the method predicts a classification *ĉ*_*i*_(*t*) (for example, if *t*_*i*_ is the test statistic, the prediction could be *ĉ*_*i*_(*t*) = 1 if *t*_*i*_ ≥ *t*, and *ĉ*_*i*_(*t*) = 0 otherwise). Across all loci, the number of true positives (TP), false positives (FP) and false negatives (FN) at the given threshold *t* is given by

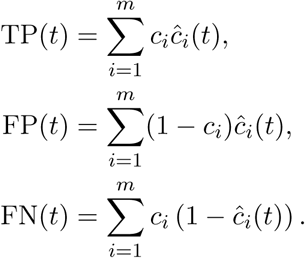

Precision and recall at this threshold are given by

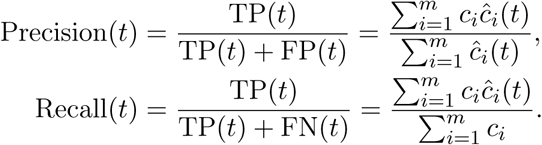

The precision-recall curve results from calculating the above two values at every threshold *t*, tracing a curve as recall goes from zero (everything is classified as null) to one (everything is classified as alternative), and the area under this curve is our final measure AUC_PR_. A method obtains the maximum AUC_PR_ = 1 if there is some threshold that classifies all loci perfectly. In contrast, a method that classifies at random (for example, *ĉ*_*i*_(*t*) ∼ Bernoulli(*p*) for any *p*) has an expected precision (= AUC_PR_) approximately equal to the overall proportion of alternative cases: 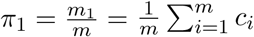. The AUC_PR_ was calculated using the R package PRROC, which computes the area by integrating the correct non-linear piecewise function when interpolating between points (Grau et al., 2015).

## 3 Results

We simulate genotype matrices and traits to go with the genotypes, in order to control important features of the population structure and to test all methods in an ideal setting where the true causal loci are known. Our simulations permit exact identification of true positives, false positives, and false negatives, ultimately yielding two measures of interest: RMSD_*p*_ measure null p-value uniformity and relates to the accuracy of type-I error control (smaller is better), while AUC_PR_ measures predictive power (higher is better) and serves as a proxy for statistical power when RMSD_*p*_ ≈ 0. However, the simulation of genotypes followed by simulation of the trait leads to a considerable amount of variance in the final measured RMSD_*p*_ and AUC_PR_, which are random variables. For that reason, every evaluation was replicated at least 10 times (varies by scenario), resulting in a distribution of RMSD_*p*_ and AUC_PR_ values per method. Except when noted, each replicate consisted of a new genotype matrix drawn from the same structure model of the scenario, followed by a new simulated trait based on this genotype matrix, which included selecting new causal loci with new effect sizes.

All scenarios are based on an admixture simulation from *K* = 10 subpopulations and a resulting generalized *F*_ST_ = 0.1, which establishes the population structure. We vary the sample size (number of individuals) in order to test the extent to which PCA overfits the population structure as the number of PCs increases (*r* ∈ {0, …, 90}), particularly in comparison to the LMM. Keep in mind that the ideal choice for the number of PCs in this simulation is *r* = *K* − 1 = 9 (the rank of the data minus the rank of the intercept). Lastly, to push all methods to their limits, we evaluate them in a scenario with both admixture and a complex family structure.

First we evaluate all methods in the large sample size scenario, which has a reasonable number of individuals (*n* = 1, 000) typical for genetic association studies. In this scenario we find a clear transition around the ideal number of PCs of *r* = 9, below of which performance is poor and above of which performance is satisfactory (Fig. 1). In particular, when *r <* 9 we find the largest RMSD_*p*_ values, which indicate that p-values are highly non-uniform and would therefore result in inaccurate type-I error control. The smallest AUC_PR_ values also occur for *r <* 9, showing that not enough PCs results in loss of predictive power as well. As expected, *r* = 9 has the best performance in terms of both RMSD_*p*_ and AUC_PR_. Remarkably, as *r* is increased up to *r* = 90, there is no noticeable change in the RMSD_*p*_ distribution, and only a small decrease in AUC_PR_ compared to the optimal *r* = 9 case. The LMM performs about as well as PCA with *r* = 9 here, with small RMSD_*p*_ values (though somewhat larger than those of *r* = 9) and larger AUC_PR_ values than PCA’s with *r* = 9. Thus, in this common scenario where sample sizes are large enough, the PCA approach with enough PCs performs as well as LMM.

**Figure 1:**
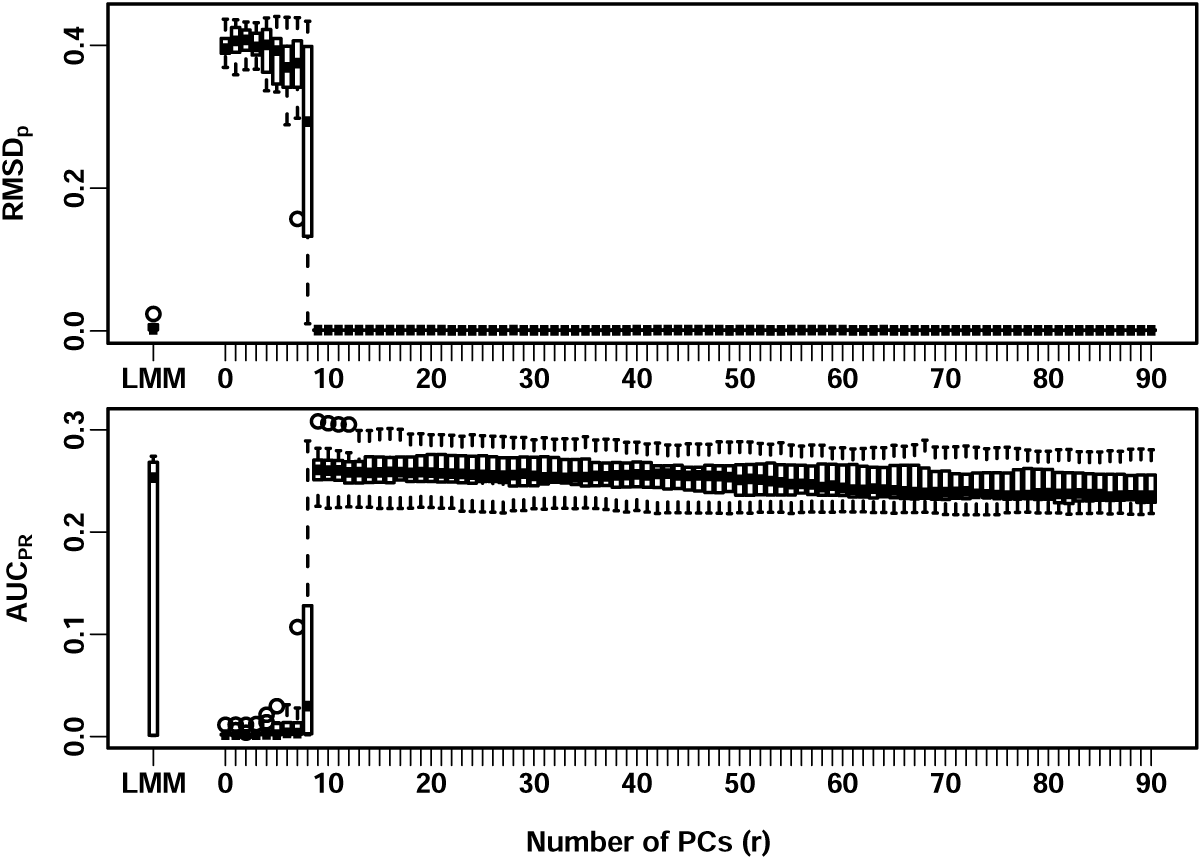
Evaluation in large sample size admixture scenario. Here there are *n* = 1, 000 individuals in the simulation. The PCA approach is tested under varying number of PCs (*r* ∈ {0, …, 90}), alongside the LMM approach (x-axis), with boxplots for 10 replicates (y-axis) for the distributions of RMSD_*p*_ (top panel) and AUC_PR_ (bottom panel). Small RMSD_*p*_ and large AUC_PR_ correspond with better performance. The ideal number of PCs is *r* = *K* − 1 = 9, where *K* is the number of subpopulations prior to admixture, which results in near zero RMSD_*p*_ and peak AUC_PR_, and performs as well as the LMM. PCA with *r <* 9 has incorrect p-values (RMSD_*p*_ ≫ 0 cases) and lowest predictive power (small AUC_PR_). Remarkably, PCA remains robust even in extreme *r >* 9 cases, with RMSD_*p*_ near zero up to *r* = 90 and minimal loss of power as *r* increases to 90.

The previous observation, that PCA continues to perform well when the number of PCs is 10 times greater than its optimum value (*r* = 90 vs *r* = 9), propelled us to find a scenario where this is no longer the case. We expect the PCA approach to begin overfitting as the number of PCs *r* approaches the sample size *n*. Increasing *r* beyond 90 does not make sense, as this would never be done in practice. Instead, we reduced *n* to 100, a number of individuals that is small for typical association studies, but which may occur in studies of rare diseases, or be due to low budgets or other constraints. To compensate for the loss of power that results from reducing the sample size, we also reduced the number of causal loci from 100 before to *m*_1_ = 10, which increases the magnitude of the effect sizes. Note that this reduction in the number of causal loci results in more discreteness in AUC_PR_ values in Fig. 2. Interestingly, we find that the relationship between RMSD_*p*_ and *r* is similar under small and large sample sizes, with ideal near-zero RMSD_*p*_ distributions for *r* ≥ 9. On the other hand, we do see a more severe overfitting effect here that results in decreased predictive power: AUC_PR_ peaks at *r* = 9 as expected, but drops more rapidly as *r* increases, with performance around *r* = 50 that is as bad as for *r* = 0, and practically zero AUC_PR_ at *r* = 90 (Fig. 2). Compared to the large sample size scenario, here LMM better matches the performance of PCA with *r* = 9.

**Figure 2:**
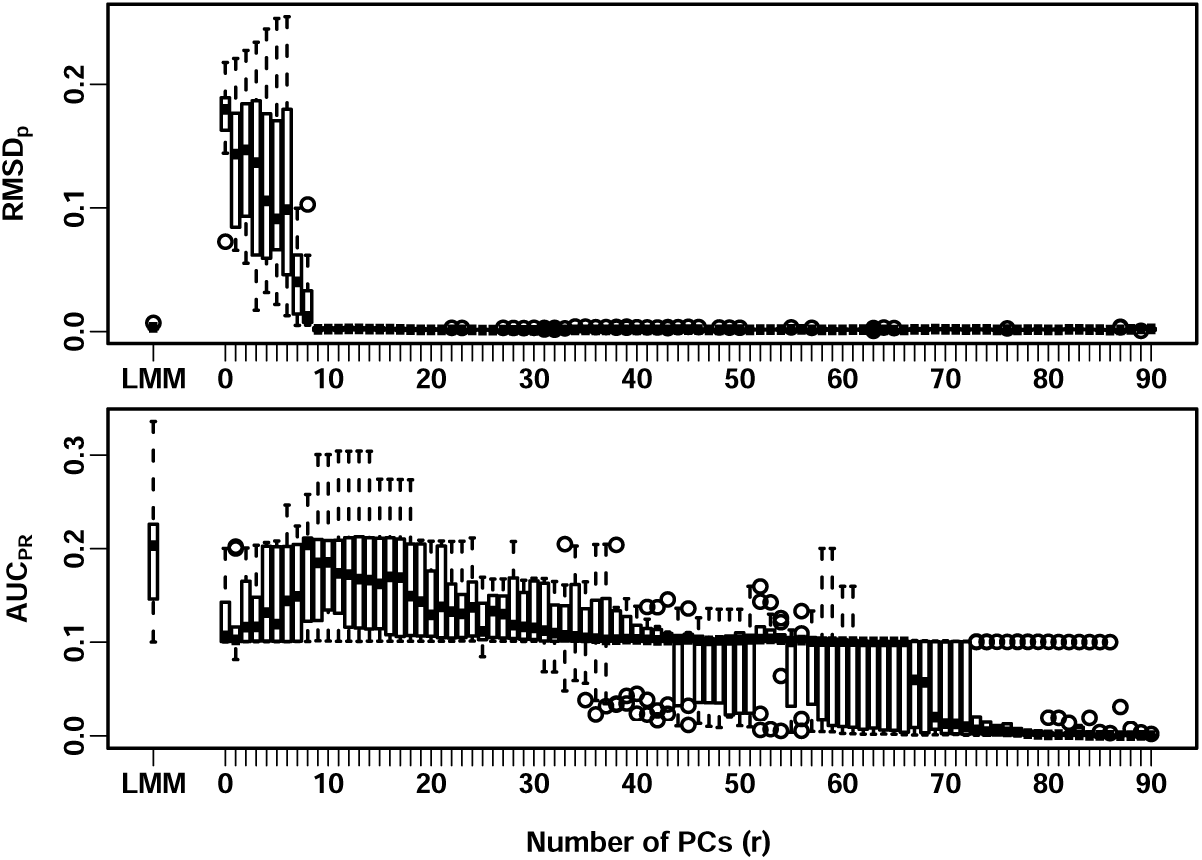
Evaluation in small sample size admixture scenario. Here there are *n* = 100 individuals in the simulation, otherwise the simulation and figure layout is the same as in Fig. 1. The pattern for RMSD_*p*_ in the top panel is similar to the previous figure. However, here there is a more pronounced drop in AUC_PR_ values as the number of PCs *r* increases from *r* = 9 to *r* = 90.

Previous work has shown that PCA performs poorly in the presence of family structure. Here we aim to characterize PCA’s behavior in a much more complex structure than before, by simulating a family of admixed founders for 20 generations, so that we may observe numerous siblings, first cousins, etc. In this case *r* = 9 is not the optimal choice, as the rank of the genotype matrix is much greater due to the family structure. We find that, although RMSD_*p*_ decreases monotonically as *r* increases, this distribution does not go to zero, instead converging to around 0.05 (Fig. 3). Additionally, the AUC_PR_ increases until *r* = 4 is reached (as opposed to *r* = 9 as before), then plateaus with marginal decreases in performance as *r* goes to 90. In contrast, the LMM does achieve a near-zero RMSD_*p*_, although the AUC_PR_ distribution is much wider than the best performing PCA cases.

**Figure 3:**
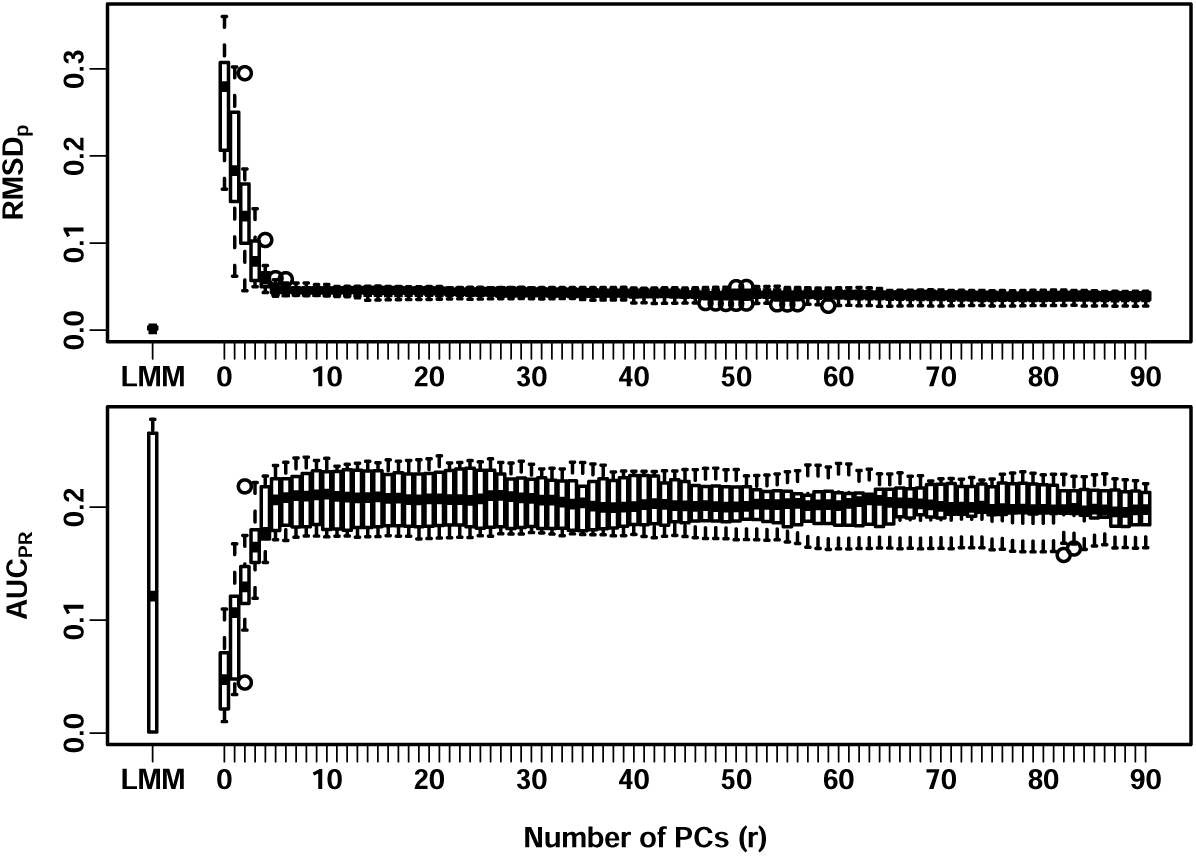
Evaluation in family structure admixture scenario. Here there are *n* = 1, 000 individuals from a family structure simulation with admixed founders and large numbers of pairs of sibling, first cousins, second cousins, etc, from a realistic random pedigree that spans 20 generations. Unlike previous figures, here RMSD_*p*_ (top panel) for PCA does not go down to zero as *r* increases. For this complex relatedness structure *r* = 9 is not the optimal number of PCs, although performance remains steady for all *r* ≥ 9 values tested.

## 4 Discussion

One important conclusion of our evaluation is that the PCA approach for genetic association studies is robust to the choice of *r* (number of PCs), as long as *r* is large enough. Thus, while we expect an *r* that is too small or too large may hurt the performance of PCA (by not modeling enough of the population structure, or by overfitting, respectively), we find that the magnitude of the performance penalty depends very strongly on whether *r* is too small or too large. In our simulations that excluded family structure, the optimal choice was *r* = *K* − 1, where *K* is the number of admixture source subpopulations, and we found that even *r* = *K* − 2 paid a large penalty in both type-I error control (measured via RMSD_*p*_) and predictive power (AUC_PR_; Figs. 1 and 2). In contrast, *r* can be much larger than its optimal value with absolutely no penalty in terms of type-I error (regardless of sample size), and only a negligible cost in predictive power when sample sizes are large (Fig. 1). The loss of predictive power by using excessive PCs is only pronounced when the number of individuals *n* is much smaller than is common nowadays, *i*.*e*. in the hundreds (Fig. 2). This robustness of PCA to the choice of *r* has long been anecdotal only (Price et al., 2006; Kang et al., 2010). We can now firmly state that it is far safer to err on the side of larger *r*, especially for large sample sizes, although testing several *r* values via simulations is always recommended.

Previous work has mostly ruled in favor of LMM instead of PCA, or otherwise tend to show comparable performance. The clearest advantage of LMM is its ability to model family relatedness. We confirm this to some extent in our family structure simulation, finding that LMM performs best in many (but not all) of our simulation replicates (Fig. 3). The other clear advantage of LMM over PCA is in having fewer degrees of freedom, which is most evident in our small sample size simulation (Fig. 2). Poor performance for PCA under small sample sizes may explain early claims that PCA was not effective in preventing test statistic inflation (Epstein et al., 2007; Kimmel et al., 2007; Luca et al., 2008).

Unexpectedly, sometimes we see LMM perform much more poorly compared to PCA in our simulation scenarios (lower quartiles in boxplots in Figs. 1 and 3). Evaluations from others also suggest that PCA can outperform a standard LMM especially when there are loci under selection or otherwise highly differentiated, and rare variants (Price et al., 2010; Wu et al., 2011; Yang et al., 2014). Thus, it seems that the additional degrees of freedom available in PCA enables it to better model loci when LMM model assumptions break. The context-dependent advantages of PCA versus LMM also argues against the simple characterization that either fixed or random effects are in principle superior models for association studies (Price et al., 2010; Sul and Eskin, 2013; Price et al., 2013; Sul et al., 2018). This reasoning also suggests that a model with both fixed and random effects (an LMM with PCs) may inherit the best of both worlds, assuming sample sizes are large enough to prevent overfitting, as some previous evaluations found (Zhao et al., 2007; Price et al., 2010).

